# Quantitative proteomics and lipidomics of TFG-deficient B cells provide insights into mechanisms of autophagic flux and plasma cell biology

**DOI:** 10.1101/2022.09.01.506221

**Authors:** Tobit D. Steinmetz, Lena Reimann, Sebastian R. Schulz, Sophia Urbanczyk, Jana Thomas, Ann-Kathrin Himmelreich, Florian Golombek, Kathrin Castiglione, Susanne Brodesser, Bettina Warscheid, Dirk Mielenz

**Author notes:** correspondence: Dr. Dirk Mielenz, Division of Molecular Immunology, Universitätsklinikum Erlangen, Nikolaus-Fiebiger-Zentrum, FAU Erlangen-Nürnberg, Erlangen, Germany, E-Mail.

## Abstract

The autophagy-flux-promoting protein TFG (*Trk-fused gene*) is up-regulated during B cell differentiation into plasma cells and supports survival of CH12 B cells. We hypothesized that quantitative proteomics analysis of CH12*tfg*KO B cells with intact or blocked autophagy-lysosome flux (via NH_4_Cl) will identify mechanisms of TFG-dependent autophagy, plasma cell biology and B cell survival. Analysis of CH12WT B cells in the presence of NH_4_Cl will identify proteins whose presence is continuously regulated by lysosomes independent of TFG. We determined hundreds of proteins to be controlled by TFG and/or NH_4_Cl. Notably, NH_4_Cl treatment alone increased the abundance of a cluster of cytosolic and mitochondrial translational proteins while it also reduced a number of proteins. Within the B cell relevant protein pool, BCL10 was reduced, while JCHAIN was increased in CH12*tfg*KO B cells. Furthermore, TFG regulated the abundance of transcription factors, such as JUNB, metabolic enzymes, such as the short-chain fatty acid activating enzyme ACOT9 or the glycolytic enzyme ALDOC. Gene ontology enrichment analysis revealed that TFG-regulated proteins localized to mitochondria and membrane-bounded organelles. Due to these findings we performed shotgun lipidomics of glycerophospholipids, uncovering that a particular phosphatidylethanolamine (PE) species, 32:0 PE, which lipidates LC3 most efficiently, was less abundant while phosphatidylglycerol (PG) was more abundant in CH12*tfg*KO B cells. In line with the role of PG as precursor for Cardiolipin (CL), the CL content was higher in CH12*tfg*KO B cells and addition of PG liposomes to B cells increased the amount of CL. We propose a role for TFG in B cell activation and plasma cell biology via regulation of proteins involved in germinal center and plasma cell development, such as BCL10 or JCHAIN, as well as in lipid homeostasis, mitochondria and metabolism.

## Introduction

During their differentiation into antibody (Ab) secreting plasma cells, B cells undergo a plethora of transcriptional, metabolic and morphologic changes (Schuh et al., 2020). These events support survival, Ab synthesis, assembly and secretion. Dimerization of IgA and pentamerization of IgM require the IgJ chain (JCHAIN). Its mRNA is expressed in all plasma cells, but JCHAIN abundance is also controlled by protein degradation (Mosmann et al., 1978). The underlying mechanism is not known (Castro and Flajnik, 2014). The antigen-specific activation of B cells depends on the BCR (Reth, 1991). After T-cell-dependent B cell activation, B cells form germinal centers to undergo positive selection, differentiation into memory B cells and plasma cells, class switch recombination and somatic hypermutation of their Ig genes by integrating signals from the BCR and CD40. These processes are coordinated by transcription factors such as nuclear factor κB (NFKB), BCL6, ID3 or JUNB (Gloury et al., 2016). Classical NFKB activation after BCR stimulation involves phosphorylation of NFKB inhibitor α (IKBA) through Inhibitor of IKBA kinase (IKK) α and β activation via IKKγ, leading to proteasomal degradation of IKBA (May and Ghosh, 1997; Thome et al., 2010). Proximal BCR signals are connected to the IKK complex and JUN N-terminal kinase (JNK) by CARMA1/CARD11 (Jun et al., 2003). CARD11 forms a complex with BCL10 and MALT1, the CBM complex. CBM null mutations abrogate antigen receptor-induced activation of NFKB, JNK and MTORC1 (Lu et al., 2018). Conversely, somatic gain-of-function mutations can cause diffuse large B cell lymphoma (DLBCL), T cell leukemia or lymphoproliferative disorders (Lu et al., 2018). Phosphorylation and ubiquitination of BCL10 can positively and negatively regulate the function and destruction of CBM complexes (Gehring et al., 2018). Moreover, the CBM complex is part of a supramolecular protein oncogenic complex involving MTORC1 and lysosomes (Phelan et al., 2018).

The JNK and MTORC1 pathways regulate autophagy (Sridharan et al., 2011) and BCL10 is a selective autophagy substrate in T cells (Paul et al., 2012). Macroautophagy/autophagy is a process starting with autophagy receptors binding ubiquitinated proteins or damaged organelles that leads to their engulfment by a double membrane vesicle. This establishes an autophagosome, which fuses with lysosomes to form an autolysosome (reviewed in (Yang and Klionsky, 2010). Engulfed cargo and their receptors become degraded during this process, which can be blocked by prevention of lysosomal acidification, for instance using NH_4_Cl (Klionsky et al., 2016). Autophagy involves a dynamic series of vesicular events managed by *ATG* (autophagy-related) genes. An important kinase controlling autophagy is ULK1 (homologous to ATG1 in yeast) (Tsukada and Ohsumi, 1993). The autophagy vesicle matures by lipidation of an ATG protein (Atg8 in yeast, LC3 and GABARAP subfamilies in mammals), which is required for autophagy vesicle expansion and closure. Plasma cells and germinal center B cells rely on autophagy (Martinez-Martin et al., 2017). In particular, autophagy cooperates with the ubiquitin-proteasome system (UPS), and ER-associated degradation (ERAD) in controlling survival of normal and malignant plasma cells (Manni et al., 2020).

In a proteomics screen for interaction partners of CARD11 we identified a 50-kDa protein named TFG (TRK-fused gene) in the mouse B lymphoma line CH27 (Grohmann et al., 2019). TFG consists of 397 amino acids structured into an N-terminal PB1 protein-protein-interaction domain (Sumimoto et al., 2007), a coiled-coil oligomerization domain and a C-terminal proline-rich region. TFG interacts with IKKγ (Miranda et al., 2006), and is involved in the structural organization of the ER (Johnson et al., 2015) as well as the ER-Golgi-intermediate compartment (ERGIC;(Hanna et al., 2017)). TFG is a regulator of the ubiquitin-proteasome system (UPS) and modifies proteasomal targeting of ER proteins (Yagi et al., 2016). While fusions of the *tfg* gene with other genes can lead to oncogenic fusion proteins that drive lymphomas and leukemias (Witte et al., 2011), *tfg* mutations can cause spastic paraplegia, optic atrophy, neuropathy and dominant axonal Charcot-Marie-Tooth disease type 2 (Nicolau and Liewluck, 2019).

We found that *Tfg* mRNA is strongly expressed in primary plasma cells compared to non-activated B cells and that TFG protein abundance is higher in *in vitro* differentiated plasmablasts than in resting B cells TFG supports survival of mouse CH12 B lymphoma cells by preventing apoptosis, by maintaining ER integrity as well as autophagy flux (Steinmetz et al., 2020). The latter has been confirmed in other cell types (Carinci et al., 2021). However, TFG is not required for induction of autophagy in B cells (Steinmetz et al., 2020). Because TFG abundance maintains the integrity of the ER and autophagy flux, regulates the UPS system and, thereby, ULK1 stability (Shi et al., 2022), we reasoned that the protein composition of CH12*tfg*WT and KO B cells should be different. Identifying changes in the proteome of CH12*tfg*WT versus KO B cells in the presence or absence of NH_4_Cl should first, reveal potential mechanisms regulating TFG-dependent survival, ER integrity and autophagy flux, which are crucial for activated B cells and plasma cells, and second, also reveal proteins whose abundance is controlled by lysosomal acidification independently of TFG.

## Results

### Identification of proteins controlled quantitatively by TFG and NH_4_Cl

Autophagy flux requires the fusion of autophagosomes with lysosomes (Klionsky et al., 2012), but can be blocked by NH_4_Cl, which prevents lysosomal function specifically (Seglen, 1975). TFG is required for autophagy flux (Carinci et al., 2021; Steinmetz et al., 2020). Proteins being degraded by autophagy flux via TFG, or being required for autophagy flux elicited by TFG, should be identified by comparing CH12*tfg*WT and KO B cells following blockage of lysosomal flux with NH_4_Cl treatment. Hence, we performed label-free quantitative proteomics analysis of CH12*tfg*WT and KO B cell clones, either treated with PBS or NH_4_Cl. We expected several groups of proteins: First, proteins up- or down-regulated by TFG independent of lysosomes; second, proteins regulated by TFG-mediated lysosomal targeting, i.e. proteins that are simultaneously more abundant in untreated CH12*tfg*KO B cells as well as in NH_4_Cl-treated CH12*tfg*WT B cells; and third, proteins that are up-regulated independently of TFG by NH_4_Cl, presumably the largest fraction. We first corroborated the reduction of autophagy flux in CH12*tfg*KO B cells by NH_4_Cl treatment and quantification of LC3-II abundance (Figure 1A, B). Cell lysates from the same experiment were subjected to quantitative proteomics analysis. In total, we quantified 3221 proteins in CH12*tfg*WT and KO B cells (supplemental table 1), of which 52 proteins were significantly up- or down-regulated solely by TFG abundance (supplemental table 2; group 1). The 15 top up- and down-regulated proteins in CH12*tfg*KO B cells are shown in Figure 1C; in line with a role of TFG in autophagy, ATG4b was regulated. The previously described knockout of TFG (Steinmetz et al., 2020) was confirmed as indicated by a large negative fold-change of ~30 (Figure 1C, supplemental table 1). Because we have used CRISPR/Cas9 to delete the *tfg* gene and because we have used different clones, we cannot fully exclude that some parts of the *tfg* gene may be expressed and translated aberrantly in very low amounts in some *tfg* KO clones. However, the method underlying the ratio calculation might also lead to a detectable, but high fold-change of a protein that is absent in the CH12*tfg*KO B cells.

**Figure 1.**
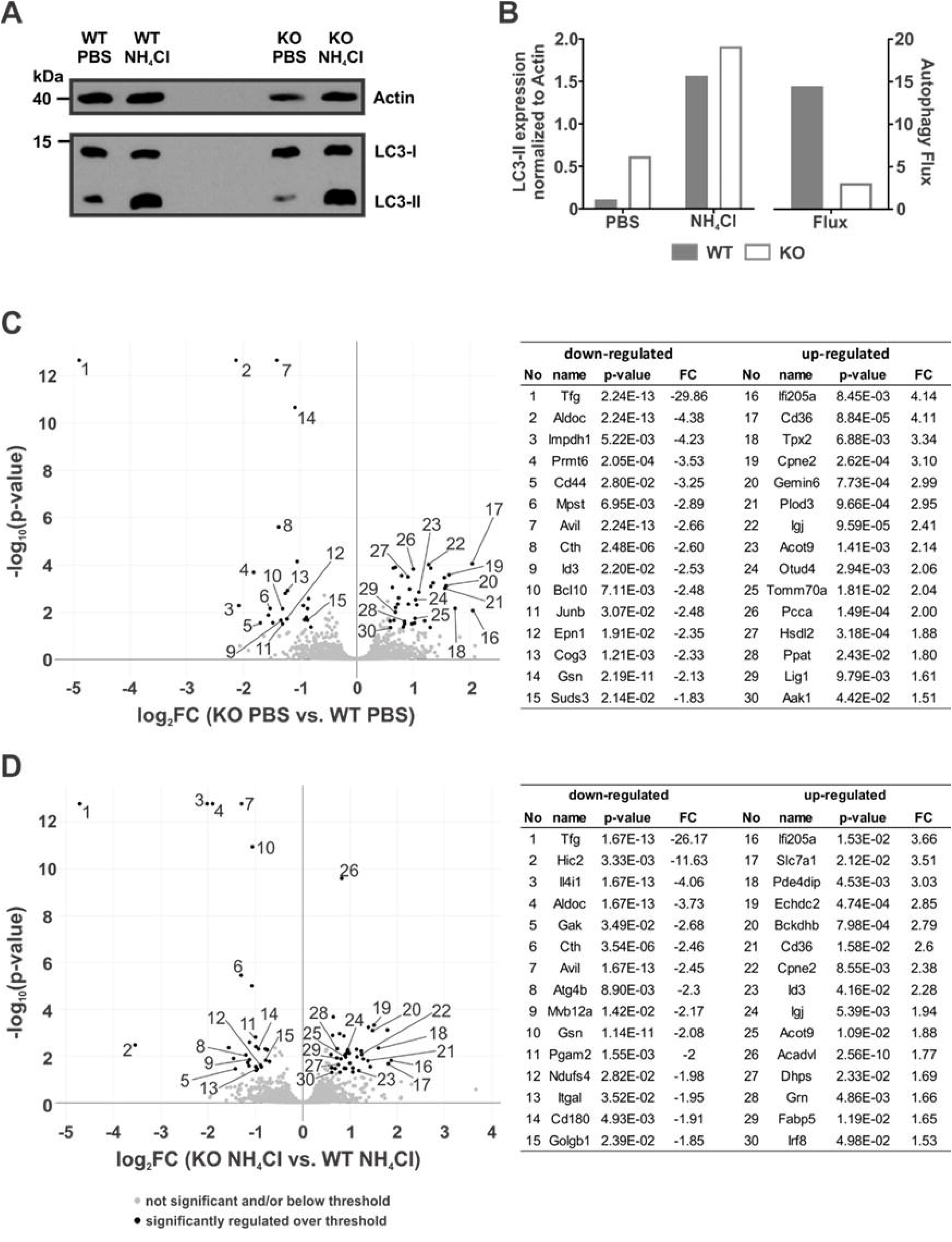
Regulation of protein abundance by TFG and NH_4_Cl in CH12 B cells. A, *CH12tfgWT* and KO B cells were left untreated or treated with NH_4_Cl for 2h. Cell lysates were separated by 15% SDS-PAGE and analyzed by Western Blotting with anti-ACTIN or LC3 antibodies. Molecular mass standards are shown on the left. B, Autophagy flux is depicted as ratio of LC3-II of NH_4_Cl treated vs. non-treated cells. C, Proteins in total cell extracts of PBS treated CH12*tfg*WT and KO B cells were digested with trypsin and analyzed by LC-MS/MS (data are combined from two experiments. In the first experiment three different *CH12tfgWT* and KO B cell clones were pooled and in the second experiment two other *CH12tfgWT* and KO B cell clones) Volcano plots and tables show the 15 most up- or down-regulated proteins in *CH12tfgWT* and KO B cells. D, Proteins in total cell extracts of NH_4_Cl treated *CH12tfgWT* and KO B cells similarly processed and depicted.

19 proteins were up-regulated in CH12*tfg*KO B cells and at the same time in CH12*tfg*WT B cells treated with NH_4_Cl (group 2) (supplemental table 2). These represent proteins that are targeted by TFG to lysosomal degradation (TPX2, GEMIN6, PLOD3, THYN1, MRRF, USF2, MRPS16, RRP9, CHTOP, PLRG1, OTUD4, TOMM70A, PCCA, DNAJC11, MRPL30, PPAT, POLD3, ETFDH, LIG1, ATAD3). Interestingly, 4 proteins were down-regulated in CH12*tfg*KO

B cells as well as in CH12*tfg*WT B cells treated with NH_4_Cl (supplemental table 2) (SUDS3, PROSER1, SH3BP1, ID3). 121 proteins were only differentially abundant in CH12*tfg*WT cells treated with NH_4_Cl compared to PBS-treated cells. 81 of those became more abundant with NH_4_Cl treatment, meaning that these proteins are delivered to the lysosomal pathway in B cells independently of TFG (group 3; supplemental table 2). Unexpectedly we also identified 42 proteins that were decreased in CH12*tfg*WT cells treated with NH_4_Cl (group 4; supplemental table 2).

### Identification of biological processes controlled by TFG and lysosomes

To reveal biological processes regulated by TFG and/or lysosomal flux, we analyzed PANTHER (http://pantherdb.org/) gene ontology (GO) processes and protein classes as well as functional clusters by STRING (https://string-db.org). Proteins that were regulated specifically by TFG but not by lysosomes (group 1; supplemental table 2) enrich for GO:0065007 (biological regulation), GO:0009987 (cellular process), GO:0051179 (localization), GO:0008152 (metabolic process) and GO:0050896 (response to stimulus). The proteins comprised the protein classes RNA metabolism, cytoskeletal proteins, transcriptional regulators, membrane traffic proteins, metabolite interconversion enzyme and protein modifying enzymes (Figure S1). Notably, the largest group contained metabolite interconversion enzymes such as ALDOC, HSDL2, ACOT9, CTH, IMPDH1 and MPST1, suggesting that TFG controls metabolic pathways via enzyme abundance. The modulation of enzyme abundance via TFG could be direct or indirect, via regulation of transcription factors such as JunB or ID3 (Figure 1C).

The 23 proteins of group 2 are of special interest because they are controlled via TFG-mediated lysosomal activity (supplemental table 2). They may represent effectors or targets of TFG-supported autophagy flux. They enrich for GO:0065007 (biological regulation), GO:0009987 (biological process), GO:008152 (metabolic process), GO:0050896 (response to stimulus) and GO:0023052 (signaling), belonging to protein classes controlling DNA metabolism, metabolite interconversion enzyme, protein modifying enzyme and translational proteins (Figure S2). STRING analysis revealed clusters of proteins involved in mitochondrial network organization (DNAJC11, ATAD3A) and mitochondrial translation (MRPS16, MRPL30, MRRF) (Figure S2).

Group 3 is composed of proteins that become more abundant with NH_4_Cl treatment independent of TFG (supplemental table 2). They are enriched in GO:0065007 (biological regulation), GO:0051179 (localization) and GO:0050896 (metabolic process), falling mainly into the classes RNA metabolism (ZNF622, SRP14, DDX1, TRMT1, RCL1, DBR1, FBL, FAM98B, ILF3, TRMT1, POLR1E, DUS3L, HDLBP), metabolite interconversion enzymes (ACLY, PFKM, ALDH18A1, ACP6, UQCR10, UQCRQ, GNPDA2, PGAM2, GPI, PRPSAP2, GMDS, NDUFV1, ELAC2, PGAM5, BLVRA), protein modifying enzyme (PITRM1, SCYL1, MAPK3, ILK, OGT) and, most robustly, a cluster of cytosolic and mitochondrial translational proteins (MRPL27, GTPBP1, MRPL3, RPL35A, RPS16, RPL34, WARS, RPS5, RPS11, FARSA, MRPL45, FARSB, MRPS2, EEFSEC, EIF2B4, RPS2, MRPL16, RPS25) (Figure S3 and S4).

Next, we conducted a PANTHER overrepresentation test, revealing that the 62 proteins regulated by TFG in total (i.e., lysosome dependent and independent), are overrepresented for the mitochondrial matrix (Figure S5).

We conclude that TFG controls the abundance of some proteins (depicted in figure 1C: ALDOC, JCHAIN, ACOT9; BCL10; see also supplemental table 2) independent of lysosomal acidification. In addition, there is an interplay between TFG and the lysosomal system that either stabilizes or destabilizes selected proteins. Notably, TFG and lysosomal inhibition seem to regulate in particular mitochondrial physiology.

### Independent verification of TFG-controlled proteins

Next, we verified the proteomics data by two independent methods, flow cytometry and Western Blot. We first focused on flow cytometry (Figure 2), commencing with JCHAIN. JCHAIN is required for dimerization and mucosal transport of IgA and for pentamerization of IgM (Castro and Flajnik, 2014). JCHAIN protein quantity is regulated on mRNA and protein level but the latter mechanism is unknown (Mosmann et al., 1978). We first confirmed that JCHAIN abundance is higher on protein level in plasma cells (primary mouse plasma cells and human B cell plasmacytoma, JK6L) than in B cells (Figure 2 A-C). We then corroborated that there is more JCHAIN in CH12*tfg*KO B cells (Figure 2D). Subsequently, we chose proteins whose abundance is only slightly regulated on the protein level (supplemental table 1) but that are known to play a role in B cells and are easily amenable to flow cytometric detection. CD20 suppresses plasma cell differentiation (Kläsener et al., 2021), CD74 is the MHCII invariant chain (Koch et al., 1991) and CD98 is an amino acid transporter and plasma cell marker (Schuh et al., 2020). Analysis of CD20, CD74 and CD98 expression by flow cytometry (Figure 2E) showed remarkably the very same pattern of regulation as with our unbiased proteomics analysis (Figure 2F). This experiment highlights the accuracy of the proteomics data. Additionally, we wished to confirm metabolic interconversion enzymes (supplemental Figure 1, also depicted in figure 1C: ALDOC, ACOT9) and BCL10, an important regulator of NFKB signaling in B cells. The relative changes of proteins observed by Western Blot did not match the proteomics data as exactly as the flow cytometry data (Figure 2F), which might be due to the superior fluorescence-based quantitative aspect of flow cytometry. Notwithstanding, the Western Blots of CH12*tfg*KO and WT B cells revealed in principle the same regulation as uncovered by proteomics: ALDOC and BCL10 were less abundant in CH12*tfg*KO B cells while ACOT9 was more abundant (Figure 2G, H).

**Figure 2.**
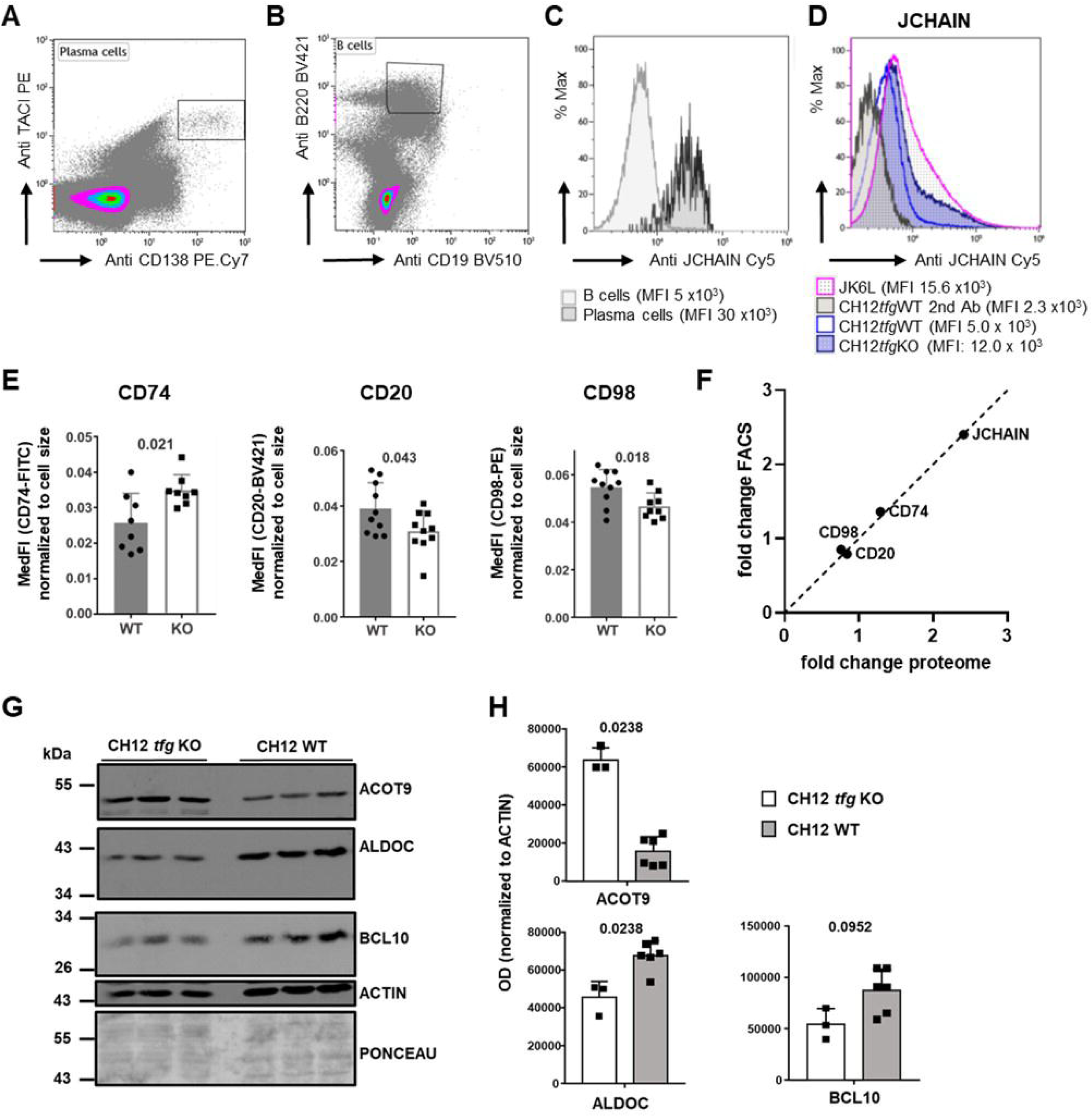
Confirmation of proteomics data by flow cytometry and Western Blotting. A-C, Mouse bone marrow cells were stained with antibodies against CD19, B220, CD138, TACI and rabbit anti JCHAIN antibody followed by a secondary anti-rabbit antibody. A, plasma cell gate, B, B cell gate, C, JCHAIN expression in B cells or plasma cells, mean fluorescence intensity (MFI) is depicted below. D, Human JK6L cells and CH12*tfg*KO and WT cells were stained with anti-rabbit antibody alone or with rabbit anti-JCHAIN antibody followed by secondary antibody. MFIs are depicted below. E, CH12tfgWT and KO B cells (7 clones each) were stained with antibodies against surface CD20 and CD98 or fixed, permeabilized and stained with antibodies against CD74 and then analyzed by flow cytometry. The fluorescence intensity was normalized to cell size and is depicted as median intensity. Each dot represents one clone with mean +/− SD. Statistical significances were calculated by Mann-Whitney U-test. F, The fold change of protein abundance of CH12*tfg*KO B cells was calculated relative to CH12*tfg*WT cells for flow cytometry and correlated to the quantification by LC-MS/MS. G, CH12*tfg*KO and WT B cells were lysed and subjected to Western Blotting with antibodies as indicated, a part of the Blot is shown. H, Quantification of the Western Blot shown in G. Background was subtracted and optical density (ODs) of the given protein were normalized to the OD of actin, Mann-Whitney U-test.

### Involvement of TFG in lipid homeostasis

Autophagy is an inherent membrane- and lipid-dependent process. B cell differentiation into plasma cells does depend on de novo lipogenesis via glucose (Dufort et al., 2014). TFG has previously been described as “lipogenic regulator” in sebocytes due to its upregulation by insulin-like growth factor (IGF)-1, leading to increased triglycerides (Choi et al., 2019). However, a clear mechanism and specific lipid classes regulated by TFG are unknown. The metabolite interconversion enzymes ACOT9 (Acyl CoA-thioesterase 9) and HSDL2 (hydroxysteroid dehydrogenase-like 2) are negatively regulated by TFG (Figure 1). ACOT9 activates short-chain fatty acids (SCFA) for de novo lipogenesis, thereby, promoting triglycerides (Steensels et al., 2020). HSDL2 is localized in peroxisomes and is presumably involved in fatty acid metabolism (Edqvist and Blomqvist, 2006). Together, these notions prompted us to address whether TFG indeed contributes to lipid homeostasis in B cells. To this end, we performed shotgun lipidomic analyses of glycerophospholipids (GPL) (Kumar et al., 2015; Urbanczyk et al., 2022) from untreated *CH12tfgWT* and KO B cells. Figure 3A depicts a representative neutral loss mass spectrum showing phosphatidylglycerol (PG) from CH12*tfg*WT and KO B cells. Notably, amongst the examined GPL, only PG was elevated in total in CH12*tfg*KO B cells, while there was no difference for total phosphatidylcholine (PC), phosphatidylethanolamine (PE), phosphatidic acid (PA), phosphatidylserine (PS) or phosphoinositols (PI) (Figure 3B). Of note, despite the total amounts of PA, PE, PI, PC and PS being similar, some particular FA species did reveal quantitative differences in CH12*tfg*KO B cells (Figure 3C-H). For instance, 32:0 PA and 32:0 PE were reduced in CH12*tfg*KO B cells while 32:0 PG and 32:0 PC were elevated. 36:2 PI was lower but 36:2 PG was higher. Some longer and unsaturated lipids, such as 38:4 PE and PI, or 40:4 PS, were also elevated.

**Figure 3.**
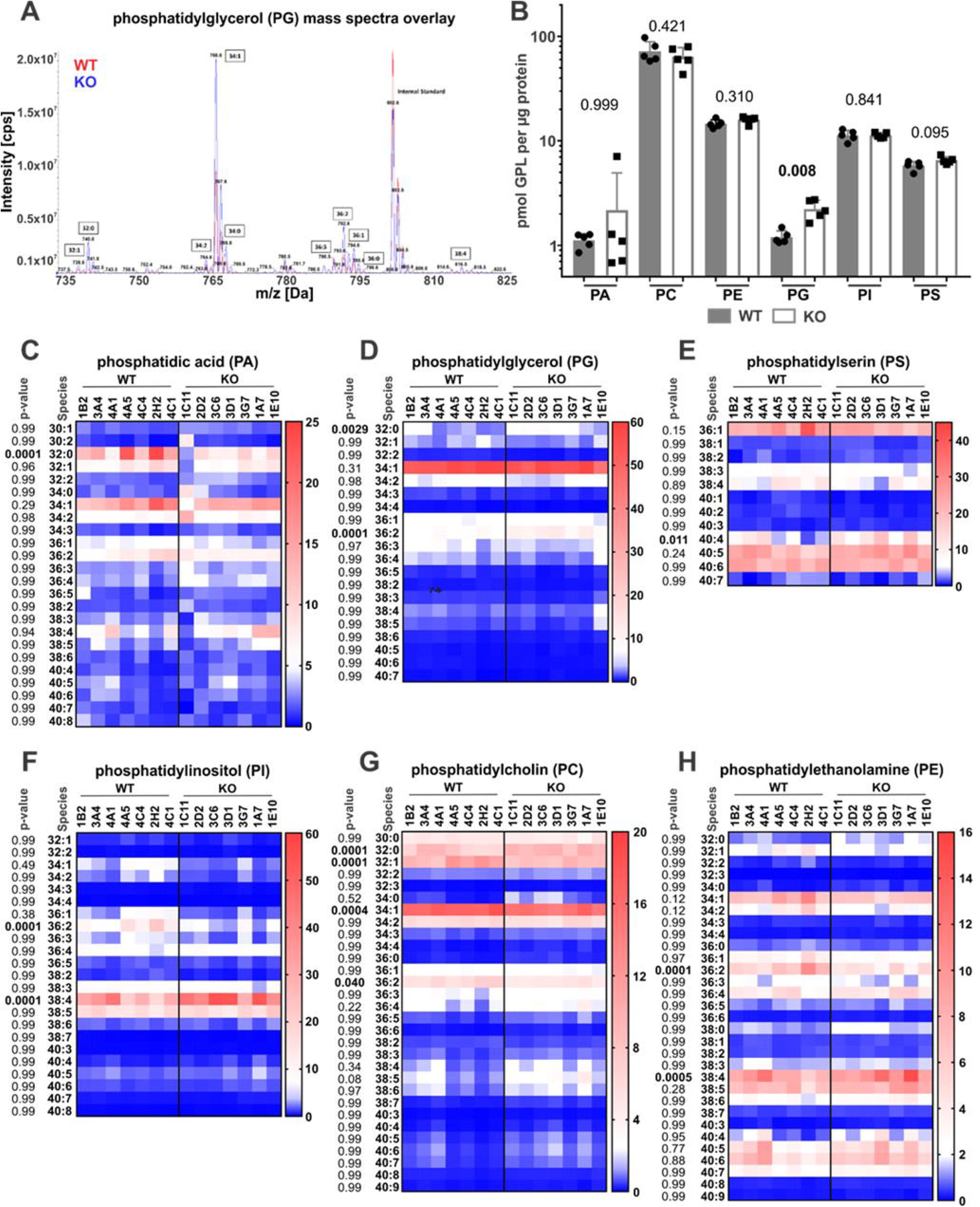
TFG limits the quantity of phosphatidylglycerol but stabilizes selected saturated glycerophospholipids in CH12 B cells. A, Glycerophospholipids of *CH12tfgWT* and KO B cells (experiment 1, 2 clones, experiment 2, 5 clones each) were analyzed via direct infusion MS/MS (*Shotgun Lipidomics*). A representative neutral loss mass spectrum of phosphatidylglycerol species from WT (red) and KO (blue) CH12 B cells (overlay) is shown, with small numbers indicating masses of the [M+NH_4_]^+^ precursor ions and large numbers indicating the total number of carbon atoms and the total number of double bonds within the two fatty acyl chains. B, The absolute abundance (pmol/μg protein) of the glycerophospholipids (GPL) phosphatidic acid (PA), phosphatidylglycerol (PG), phosphatidylinositol (PI), phosphatidylcholine (PC), phosphatidylethanolamine (PE) and phosphatidylserine (PS) in *CH12tfgWT* and KO B cells is depicted (mean −/+ SD, each dot represents on clone). Significance was calculated using Mann-Whitney U-test. C-H, The relative abundance (Mol%) of subspecies of phosphatidic acid (PA), phosphatidylglycerol (PG), phosphatidylinositol (PI), phosphatidylcholine (PC), phosphatidylethanolamine (PE) and phosphatidylserine (PS) glycerophospholipids of *CH12tfgWT* and KO B cells is depicted as heatmap. Each symbol represents one clone. P-value and lipid species on the left of each heatmap. The first number depicts the total number of carbon atoms, the second number the total number of double bonds within the two fatty acyl chains. Combined results from 2 experiments, N=2, n=7; statistics calculated using 2-way ANOVA

### Functional consequences of altered PG in CH12*tfg*KO B cells

TFG appears to control the abundance of proteins localized in the mitochondrial matrix and in the mitochondrion (Figure S5). PG is the precursor for Cardiolipin (CL), a specific and important lipid of the inner mitochondrial membrane (IMM) (Athenstaedt and Daum, 1999; Short and White, 1972). We found PG up-regulated in CH12*tfg*KO B cells (Figure 3). To test whether the elevated PG content in CH12*tfg*KO B cells influences CL abundance we quantified CL by flow cytometry with 10-N-Nonyl Acridine Orange (Mileykovskaya et al., 2001) (Figure 4A). We found a trend towards increased CL in CH12*tfg*KO B cells and speculated that there is a steady-state flux of endogenous PG towards CL, which is then further metabolized. Therefore, we created an acute situation by adding 32:0 PG liposomes to B cells. We chose primary, short-term activated B cells to have a more physiologic setting. In fact, exogenous addition of PG elevated CL. We propose that TFG, by modulating the abundance of lipid modifying enzymes, controls lipid homeostasis, in particular PG and CL, impacting thereby presumably on mitochondrial metabolism.

**Figure 4.**
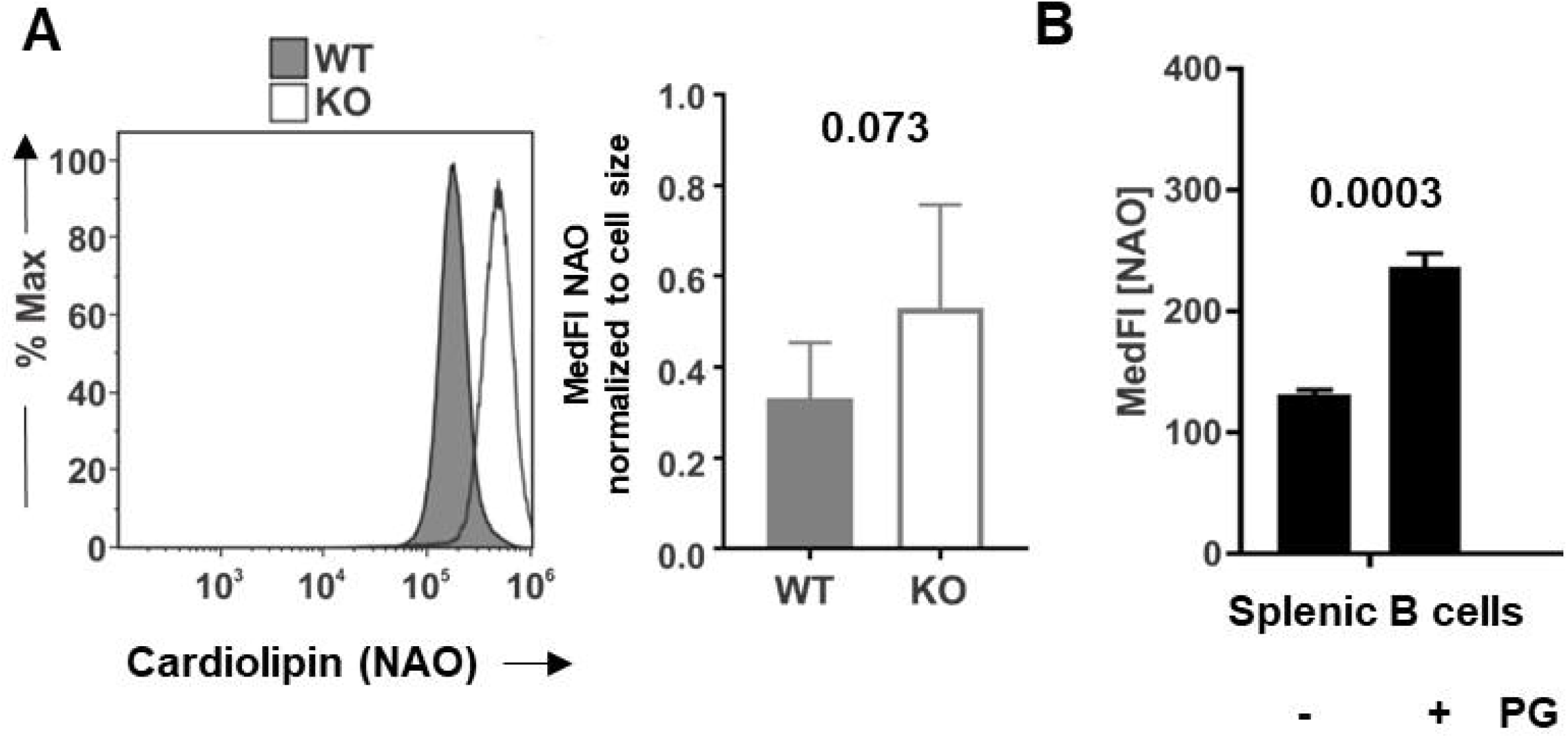
TFG and phosphatidylglycerol dependent effect on cardiolipin. A, CH12*tfg*WT and KO B cells were stained with 10-N-Nonyl acridine orange (NAO) and analyzed by flow cytometry. The left panel shows a representative histogram, the right panel depicts the median fluorescence intensity normalized to cell size (mean −/+ SD, 7 clones each). B, Phosphatidylglycerol liposomes (PG 36:2) were added to murine splenic B cells activated with LPS for 24h. Cells were stained with NAO and analyzed by flow cytometry. The median fluorescence is depicted as mean −/+ SEM, statistics with paired t-test.

## Discussion

Autophagy is crucial for humoral immunity but there are few characterized regulators apart from the canonical ATG genes in B cells. To gain more insights into mechanisms and consequences of autophagy we characterized the proteome of murine CH12*tfg*KO and WT B lymphoma cells in steady state or when lysosomal activity was inhibited by NH_4_Cl. In total, we identified 3221 proteins in CH12 B cells. Even changes in protein abundances below 1.5-fold between treatment and control were validated by complementary methods, such as flow cytometry, attesting to the quantitative aspect of the proteomic data. The quantification of protein abundance in the absence or presence of NH_4_Cl or TFG identified clusters of proteins that were either regulated by NH_4_Cl treatment, TFG (i.e. genotype-driven) or both. By this approach, we identified substrates and processes, such as lipid homeostasis, downstream of TFG and lysosomal activity. Importantly, we also identified proteins regulated by lysosomes independently of TFG. Interestingly, many proteins that appear to be stabilized by NH_4_Cl, i.e. that are targeted to Iysosomes, are proteins of the cytosolic and mitochondrial translation machinery. It is tempting to speculate that lysosomal degradation aids the catabolic function of autophagy by limiting protein translation, consistent with a positive regulation of autophagy by TOR inhibition (Blommaart et al., 1995; Noda and Ohsumi, 1998). In general, our data will help to understand autophagy and lysosome-controlled processes, not only in B cells. To our surprise, a large fraction of proteins was not up-, but down-regulated by NH_4_Cl. Because lysosomal and UPS pathways are intertwined (Korolchuk et al., 2010), the proteasome may take over under conditions of lysosomal inhibition.

Along these lines, proteins regulated by TFG, but not by NH_4_Cl, could well be linked to the function of TFG in the UPS system (Yagi et al., 2016), a possibility that warrants further investigation. Interestingly, ATG4B was down-regulated in CH12*tfg*KO B cells treated with NH_4_Cl. ATG4B is a protease that immediately removes the C-terminal residues of LC3 to produce the LC3-I isoform (Yang et al., 2021). During autophagy flux, the LC3-I isoform is converted to LC3-II by the addition of PE to the new C-terminus. However, inactive ATG4B blocks lipidation of LC3 paralogues, resulting in the inhibition of autophagy (Yang et al., 2021). Thus, the regulation of ATG4B abundance by TFG and NH_4_Cl provides a mechanism of TFG-mediated autophagy flux. Moreover, LC3 lipidation works best with 36:2 PE (DOPE)(Nath et al., 2014). Strikingly, exactly this particular PE species was reduced in CH12*tfg*KO B cells that exhibit reduced autophagy flux. Hence, the combined regulation of ATG4B and 36:2 PE by TFG may underly autophagy flux in CH12 B cells.

GSN (Gelsolin) was down-regulated, which is a sign of ER stress and results in EIF2A phosphorylation(Roy et al., 2018). Moreover, PRMT6 was also reduced which could lead to autophagy induction(Frankel et al., 2002). The regulation of these proteins is in line with our previous notion that overt ER stress is perpetuated in CH12*tfg*KO B cells because autophagy flux, an inherently lysosome-controlled process, is simultaneously reduced. Players related to ER stress could also be MPST and CTH which are less present in CH12*tfg*KO cells and regulate H_2_S and autophagy in a Redox-driven interplay (Chen et al., 2018; Panagaki et al., 2020), possibly impacting redox-sensitive ATG4 (Perez-Perez et al., 2014).

One surprising GO term characterizing TFG-regulated proteins was “mitochondrial matrix”, suggesting a possible involvement of TFG in metabolism. The finding that TOMM70A was elevated in CH12*tfg*KO B cells could point to impaired mitophagy (Bertolin et al., 2013). Albeit we did not observe signs of defective mitochondria by electron microscopy (Steinmetz et al., 2020) this may deserve further investigation and impaired mitophagy might feed back on mitochondrial function.

An intriguing result was that some of the most quantitatively TFG-suppressed proteins are ACOT9, CD36 and HSDL2, all of which modify lipid synthesis and homeostasis (Tillander et al., 2014). A previous study has shown that TFG protein was increased in a time- and dose-dependent manner by IGF-1 in sebocytes and that TFG overexpression increased triglycerides globally as assessed by thin layer chromatography (Choi et al., 2019). ACOT9 promotes triglyceride biosynthesis (Steensels et al., 2020). Consequently, we reasoned that there should be alterations in lipids in CH12*tfg*KO B cells and therefore we covered the glycerophospholipid pool of CH12*tfg*KO B cells in depth by neutral loss mass spectrometry. The main finding was the limitation of PG abundance by TFG. PG is the biosynthetic precursor of CL (Short and White, 1972). The fact that there was only a trend towards the elevation of CL by TFG deficiency may be explained by an equilibrium of CL synthesis from PG and by its turnover. This interpretation is supported firstly, by the finding that acutely added exogenous PG elevated CL, and secondly, that phosphoglycerolipids with similar acyl side chains showed a disequilibrium in CH12*tfg*KO B cells. Specifically, 32:0 PA and 32:0 PE were reduced in CH12*tfg*KO B cells while 32:0 PG and 32:0 PC were elevated; 36:2 PI was lower but 36:2 PG was higher. There may also be an ongoing conversion of 32:0 PA to 32:0 PG via lipid phosphate phosphohydrolase-driven generation of 32:0 diacylglycerol in CH12*tfg*KO B cells (Athenstaedt and Daum, 1999).

Concerning B cells, TFG controlled the abundance of proteins involved in B cell activation (BCL10) (Thome et al., 2010), the germinal center reaction (ID3), (Gloury et al., 2016), plasma cell differentiation (JCHAIN) or B cell activation and lymphomas (BCL10, JUNB). In contrast to the selective targeting of BCL10 by autophagy (Paul et al., 2012), TFG stabilized BCL10. The stabilization of BCL10 abundance by TFG is congruent with our finding of TFG as part of a CARD11 complex in B cells and suggests that TFG is a positive regulator of the CBM pathway. This is in line with its role as a positive regulator of NFKB activity (Miranda et al., 2006). ID3 becomes up-regulated by autophagy, albeit in macrophages (Sanjurjo et al., 2018), and IRF8 drives autophagy(Gupta et al., 2015). The fact that TFG elevates also JUNB protein while JUNB is a negative regulator of autophagy may constitute a feedback loop (Lin et al., 2021).

We also found suppression of JCHAIN protein abundance by TFG. The mechanism behind this phenomenon requires further investigation, which is beyond the scope of this manuscript. However, TFG may be a strong candidate for the regulation of JCHAIN in different plasma cell subsets, which is a yet unresolved but crucial issue for understanding IgA and IgM homeostasis as well as the regulation of anti-inflammatory antibody responses (Castro and Flajnik, 2014).

TFG has recently been reported to aid in the antiviral response against Sendai virus by promoting TBK1 and IRF3 activation via their transport to MAVS-containing compartments (Khan et al., 2021). Here we show that loss of TFG promotes an extensive increase of the interferon-regulated protein IFI205, which would be pro-inflammatory via regulation of the inflammasome (Ghosh et al., 2017). Along this line, OTUD4 is also up-regulated in CH12*tfg*KO B cells and one function of OTUD4 is stabilization of the antiviral protein MAVS (Liuyu et al., 2019). Therefore, TFG may have diverse pro- or anti-inflammatory functions depending on the type of infection, its expression level and the cellular context. Those possibilities can only be addressed in conditional *tfg*^-/-^ mice.

In summary, the work presented here generated new insights and hypotheses on the function of TFG in - and beyond - the regulation of ER integrity and autophagy flux, such as the regulation of inflammation, redox balance, mitochondrial function or lipid homeostasis. Testing those hypotheses will aid in understanding neurodegeneration and immunity.

## Abbreviations

Ab: antibody
Ag: antigen
ASC: antibody-secreting cells
ATG: autophagy-related
BCR: B cell receptor
COPII: coat protein complex II
CpG: non-methylated CpG oligonucleotide
ER: endoplasmic reticulum
ERAD: ER-associated degradation
FO: follicular
GO: gene ontology
HC: heavy chain
Ig: immunoglobulin
LC: light chain
NFKB: nuclear factor of kappa light polypeptide gene enhancer in B cells
TLR: toll-like receptor
UPR: unfolded protein response.

## Acknowledgements

We thank the PRIDE team for data deposition to the ProteomeXchange Consortium. This work was funded by the Deutsche Forschungsgemeinschaft (DFG; transregional collaborative research grant TRR130, to D.M. and B.W., research training grant 1660, to D.M. and Mi 939/6-1, to D.M.). S.B. was supported by the Cologne Excellence Cluster on Cellular Stress Responses in Aging-associated Diseases – CECAD). Research in the Warscheid lab is further supported by the DFG Project ID 403222702/SFB 1381 and the Germany’s Excellence Strategy (CIBSS – EXC-2189 – Project ID 390939984).

## Materials & Methods

### B Cell culture

CH12*tfg*WT and CH12*tfg*KO B cells (Steinmetz et al., 2020) were cultured in R10 medium (RPMI1640 [Gibco Life Technologies, 31870-025], 10% FCS, 2 mM glutamate [Gibco Life Technologies, 25030-024], 1 mM sodium pyruvate [Gibco Life Technologies, 11360-039], 50 U/ml penicillin G, 50 μg/ml streptomycin [Gibco Life Technologies, 15140-122], 50 μM β-mercaptoethanol [Gibco Life Technologies, 31350-010]) at 37°C and 5% CO_2_. JK6L cells were cultured in R10 medium supplemented with supernatants of IL-6-producing J558 cells (Meister et al., 2007). Primary mouse B cells were prepared form spleen and activated with lipopolysaccharide (LPS) for 24h as described (Urbanczyk et al., 2022).

### Glycerophospholipid analysis

Glycerophospholipids (PC, PE, PI, PS, PG, PA) in B cells were analyzed by Nano-Electrospray Ionization Tandem Mass Spectrometry (Nano-ESI-MS/MS) with direct infusion of the lipid extract (*Shotgun Lipidomics*): 14 to 45 × 10^6^ cells were homogenized in 300 μl of Milli-Q water using the Precellys 24 Homogenisator (Peqlab, Erlangen, Germany) at 6.500 rpm for 30 sec. The protein content of the homogenate was routinely determined using bicinchoninic acid. To 100 μl of the homogenate 400 μl of Milli-Q water, 1.875 ml of methanol/chloroform 2:1 (v/v) and internal standards (125 pmol PC 17:0-20:4, 132 pmol PE 17:0-20:4, 118 pmol PI 17:0-20:4, 131 pmol PS 17:0-20:4, 62 pmol PG 17:0/20:4, 75 pmol PA 17:0/20:4; Avanti Polar Lipids) were added. Lipid extraction and nano-ESI-MS/MS analysis were performed as previously described (Kumar et al., 2015). Endogenous glycerophospholipids were quantified by referring their peak areas to those of the internal standards. The calculated glycerophospholipid amounts were normalized to the protein content of the tissue homogenate.

### Western Blot

Cells were washed in phosphate-buffered saline (PBS) and lysed in buffer containing 1% Triton X-100, 150 mM NaCl, 25 mM Tris/HCl pH 7.5, 5 mM EDTA, 1 mM phenyl-methyl-sulfonyl-fluoride on ice for 15 min. The lysate was cleared by centrifugation at 10000g at 4° C for 15 min and supernatants were prepared for sodium dodecyl sulfate (SDS)-polyacrylamide gel electrophoresis (PAGE) according to standard procedures. After SDS-PAGE, proteins were transferred onto a nitrocellulose membrane, the membrane was washed with deionized water and stained with Ponceau S. The membrane was blocked with 5% skim milk powder in TBST (150 mM NaCl, 25 mM Tris/HCl pH 7.5, 0.1 Tween-20), followed by incubation with primary antibody (diluted in TBST containing 3 % bovine serum albumin (BSA), 0.1 % NaN_3_) and 4 washes with TBST. The membrane was incubated with secondary horseradish peroxidase (HRP) conjugated antibody diluted in 5 % skim milk powder in TBST. Anti-TFG antibody was described previously (27) (19)(3). Other antibodies were: mouse anti-ACTIN (Sigma Aldrich, A1978), rabbit anti-ACTIN, ACOT9, ALDOC (Sigma Aldrich, A2066, HPA035533, HPA003282), rabbit anti-MAP1LC3B/LC3 and BCL10 (Cell Signaling Technology, 2775S and 4237S), goat anti-rabbit IgG-HRP (Jackson Immunoresearch, 111-035-046), goat anti-mouse IgG-HRP (Jackson Immunoresearch, 115-035-008). After washing, the membrane was incubated with enhanced chemiluminescence (ECL) before exposure to X-ray films. Quantification was performed by densitometry of scanned X-ray films with ImageJ 1.52p. Band intensities were measured and the background of X-ray films was subtracted.

### Generation of tryptic peptides

Proteomic analysis was performed as described. (Gronemeyer et al., 2013). Briefly, we pooled in total 1.5 × 10^8^ cells from each 5 WT and 5 *tfg*KO subclones of CH12 cells. Clones were treated individually for each experimental condition with 1 % phosphate-buffered saline (PBS) or with 1% of 5M NH_4_Cl for 2h in complete RPMI160 medium under normal culture conditions. The samples were washed with PBS and split into technical replicates of 1×10^7^ cells. Pelleted cells were shock frozen using liquid nitrogen and stored at −80° C until further processing. Cells were lysed in 30 mM Tris base with 7 M urea and 2 M thiourea, pH 8.5. Cells were sonicated on ice twice with an ultrasonic homogenizer for 15 s at 90% of maximum power. Cell debris was removed by centrifugation for 20 min at 21.000 x g and 4°C. Samples containing 20 μg of total protein were prepared and diluted with 50 mM ammonium bicarbonate to 2M Urea and incubated for 20 min with 1 μL Benzonase at room temperature (RT). Cysteine residues were reduced with 5 mM TCEP, alkylated with 50 mM CAM for 5 min at 45 °C as described before (Reimann et al., 2020). Each sample was then digested with Lys-C (protein to enzyme ratio 1:100, FUJIFILM Wako Chemicals Europe GmbH, Neuss, Germany) for 30 min at 37°C Subsequently, trypsin (protein to enzyme ratio 1:50, Promega, Mannheim, Germany) was added for overnight digestion at 37°C. The reaction was stopped by acidification with TFA (1% [v/v] final concentration) and 10 μg of the peptide sample were desalted using an in-house prepared C18 Stage tip cartridge using 3 layers of C18 material. The cartridges were washed with 1 ml methanol followed by 1 ml 70% ACN with 0.1% TFA. The column was conditioned twice with 0.4 mL 0.1% TFA before the sample was applied. Peptides were washed with 0.4 0.1% TFA and eluted with two times 30 μL 70% ACN. Peptides were dried *in vacuo* using a rotational vacuum concentrator (Christ, Osterode am Harz, Germany).

### Proteomics analysis

Reversed-phase HPLC was performed on an UltimateTM 3000 RSLCnano system (Thermo Fisher Scientific, Dreieich, Germany) equipped with two Waters pre-columns and a Waters nano *m/z* analytical column (75 μm × 250 mm, 3 μm, 100 Å, Thermo Fisher Scientific) (Gronemeyer et al., 2013). The HPLC was connected to a QExactive Plus instrument with the following parameter settings: spray voltage, 1.8 kV; capillary voltage, 200 V; automatic gain control (AGC), 3×10^6^ ions; max. ion time (IT), 60 ms. Full scans were acquired in the orbitrap with a mass range of *m/z* 375 to 1,700 and a resolution (R) of 70,000 at *m/z 2*00. A TOP12 method was used and parameters were as follows: normalized collision energy (NCE), 28%; dynamic exclusion (DE) 45 s; AGC, 5,000; max. IT, 120 ms. A solvent system consisting of 0.1% FA (solvent A) and 86% ACN, 0.1% FA (solvent B) was used for peptide separation. The RSLC was operated with a flow rate of 0.250 μl/min for the analytical column.

Raw data were searched against the Uniprot *Mus musculus*. Proteome set using MaxQuant version 1.5.4.0. The species was restricted to MaxQuant default settings unless stated otherwise. Database searches were conducted with trypsin and Lys-C (for quantitative proteome analysis) as proteolytic enzymes, a maximum number of three missed cleavages, and mass tolerance of 4.5 ppm for precursor and 0.5 Da (CID data) or 20 ppm (HCD data) for fragment ions. Two missed cleavages were allowed. Carbamidomethylation of cysteine residues was set as fixed modification, and oxidation of methionine and N-terminal acetylation were applied as variable modifications. The minimum number of unique peptides was set to 1. A false discovery rate (FDR) of 1% was applied to both peptide and protein lists. For global proteome analysis, “label-free” and “iBAQ” quantification as well as “match between runs” were enabled with default settings.

### Bioinformatic analysis

Functional analysis of proteomics data was performed using Microsoft R studio and DEP package version 1.4.1 for single protein expression values for 3221 quantified proteins. In brief, raw expression data were adjusted to contain only protein ID, Uniprot code, protein name, gene name and expression values of the 20 samples. Raw data were imported into R studio and supplemented with meta data to assign raw data with sample information.

Duplicates were excluded within the different ID columns by making the data set unique and log2 transformation of the data set was reversed. Subsequently, data was written into a summarized experiment format and normalized. Using protein-wise linear models combined with an empirical t-test based Bayes statistics p-value calculation for the comparison pairs WT/PBS + KO/PBS, KO/PBS + KO/NH4Cl, WT/NH_4_Cl + KO/NH_4_Cl and WT/PBS + WT/NH_4_Cl was performed. Significant data were filtered using the cut-off values of p<0.05 and expression fold change >1.5. Exported data sets were used for the analysis of regulation of single protein expression. Volcano plots were generated using DEP plot. The IDs of significantly regulated proteins were functionally classified using gene ontology (GO) enrichment analysis (http://pantherdb.org/) (default settings) (Thomas et al., 2022). Functional protein clusters were identified by STRING (Mus musculus proteins, default settings) (https://string-db.org/) (Szklarczyk et al., 2021).

### Flow cytometry and cell sorting

2×10^6^-4×10^6^ cells were pelleted in FACS tubes (Micronic, Lelystad, Netherlands) at 300 x g for 5 min at 4°C and resuspended in 50 μl of unlabeled anti-FCGR3/CD16-FCGR2B/CD32 Ab (Invitrogen, 14-0161-86) 10 μg/ml in FACS-buffer (PBS, 2% fetal calf serum [FCS, Gibco Life Technologies, 10270-106], 0.05% sodium azide [Carl Roth, K305.4]) for 15 min on ice. Cells were washed once with FACS-buffer by centrifugation at 300 x g for 5 min at 4°C, resuspended in 50 μl FACS-buffer containing the respective fluorochrome-coupled Abs (anti-TACI/CD267 PE, clone eBio8F10-3, eBioscience, anti CD138 PECy7, clone 281-2, Biolegend, anti CD19 BV510, clone 6D5, Biolgegend, anti B220 BV421, clone RA3-6b2, Biolegend, anti CD98PE, #128207, Biolegend, anti-CD74-FITC, clone In-1, Pharmingen, anti-CD20-APC.Cy7, #302314, Biolegend) and incubated for 20 min on ice in the dark. Intracellular staining was performed after surface staining, fixation and permeabilization and rabbit anti JCHAIN antibody SP105 (Thermo, MA5-16419), followed by a secondary goat anti-rabbit antibody conjugated to Alexa 647 (Jackson). Cells were washed twice with FACS-buffer by centrifugation at 300 x g for 5 min at 4°C. Data were acquired using a Gallios flow cytometer (Beckman Coulter, Brea, USA). Analyses were performed using Kaluza versions 1.3 and 2.1 (Beckman Coulter, Brea, USA). For intracellular flow cytometric, the fix and perm kit from An der Grub (distributed by DIANOVA, Hamburg, Germany) was used according to the manufacturer’s instructions. Cardiolipin was detected by 10-N-nonyl acridine orange (NAO) (Sigma-Aldrich).

### Liposome production

Liposome production was based as recently described (S. T. Poschenrieder, 2016; Urbanczyk et al., 2022). Here, 11.4 mL of vesicle buffer (10 mM Tris-HCl, 150 mM NaCl, pH 8.0) were poured into unbaffled miniaturized stirred tank reactors (BioREACTOR48, 2mag, Munich, Germany) and stirred at 4000 rpm at 25°C using an S-type stirrer. Then, 600 μL of the dissolved phospholipid 16:0 PG (1,2-dipalmitoyl-sn-glycero-3-phosphate) (10 mM in chloroform) were injected under stirring, leading to a whitish highly dispersed emulsion. The reactor shell, originally made of polystyrene, has been replaced by borosilicate glass, to avoid damage by the chloroform. The process was continued, until the solution became transparent, indicating the evaporation of the chloroform, which was the case after 6 hours. Subsequently, the solution was centrifuged at 13.000 rpm in a tabletop centrifuge to remove precipitates. The liposomes in the supernatant were then concentrated to 1.5 mg mL^-1^ by centrifugation for 50 min at 50.000 g and resuspending of the resulting pellet in the appropriate amount of vesicle buffer. The quality of the liposomes during and after the production process was monitored via dynamic light scattering using a Zetasizer NS (Malvern, Worcestershire, UK) as reported.

---

**Figure S1.**
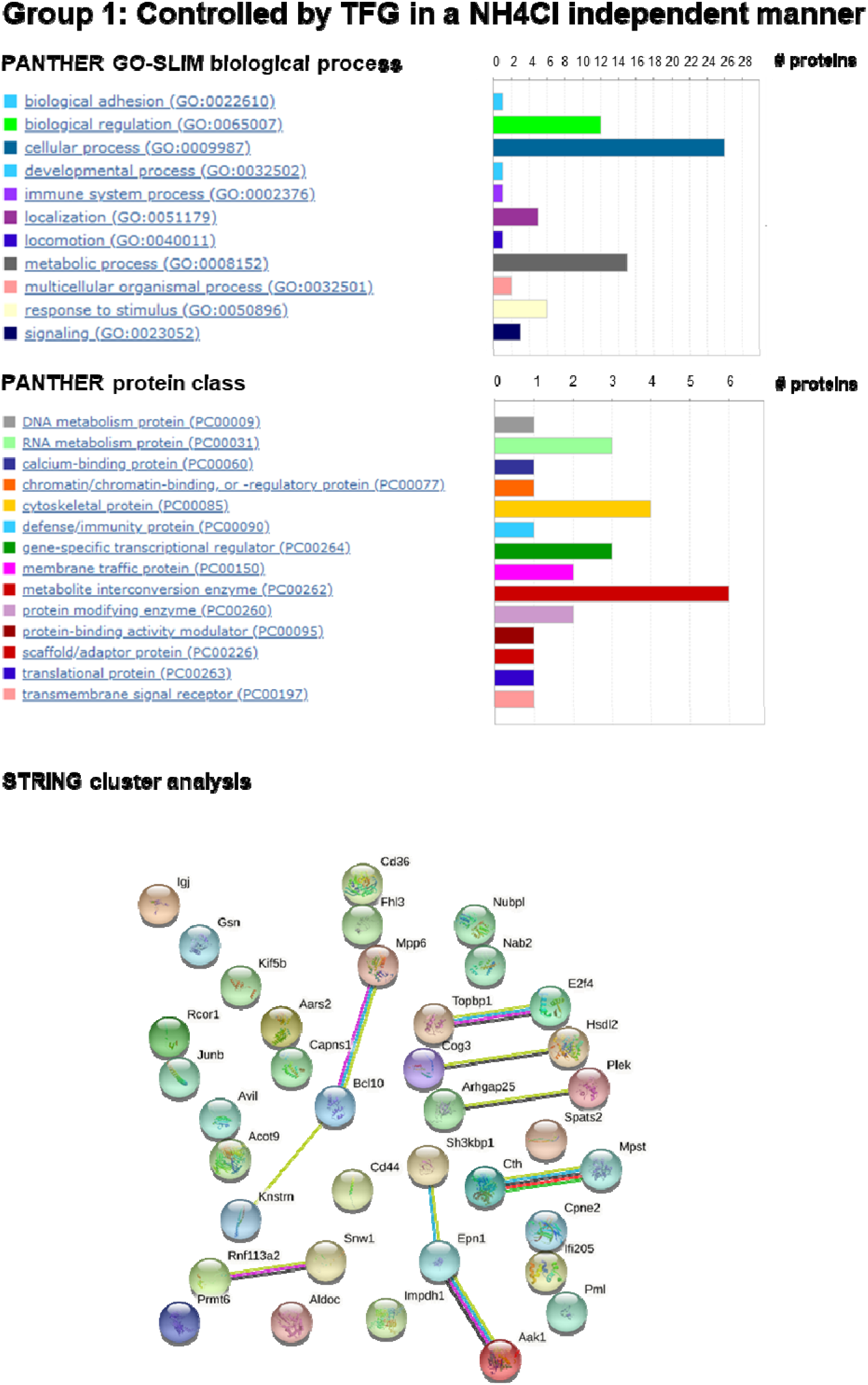
PANTHER GO-SLIM/ protein classifications and STRING analysis of TFG regulated proteins. Analyses were performed with the proteins: IFI205A, CD36, CPNE2, IGJ, ACOT9, KNSTRN, SNW1, HSDL2, KIF5B, NUBPL, AARS2, SH3KBP1, RCOR1, MPP6, SPATS2, AAK1, TOPBP1, NAB2, FHL3, PML, E2F4, CAPNS1, ARHGAP25, GSN, COG3, EPN1, BCL10, JUNB, CTH, AVIL, PLEK, MPST, RNF113A2, CD44, PRMT6, IMPDH1, ALDOC.

**Figure S2.**
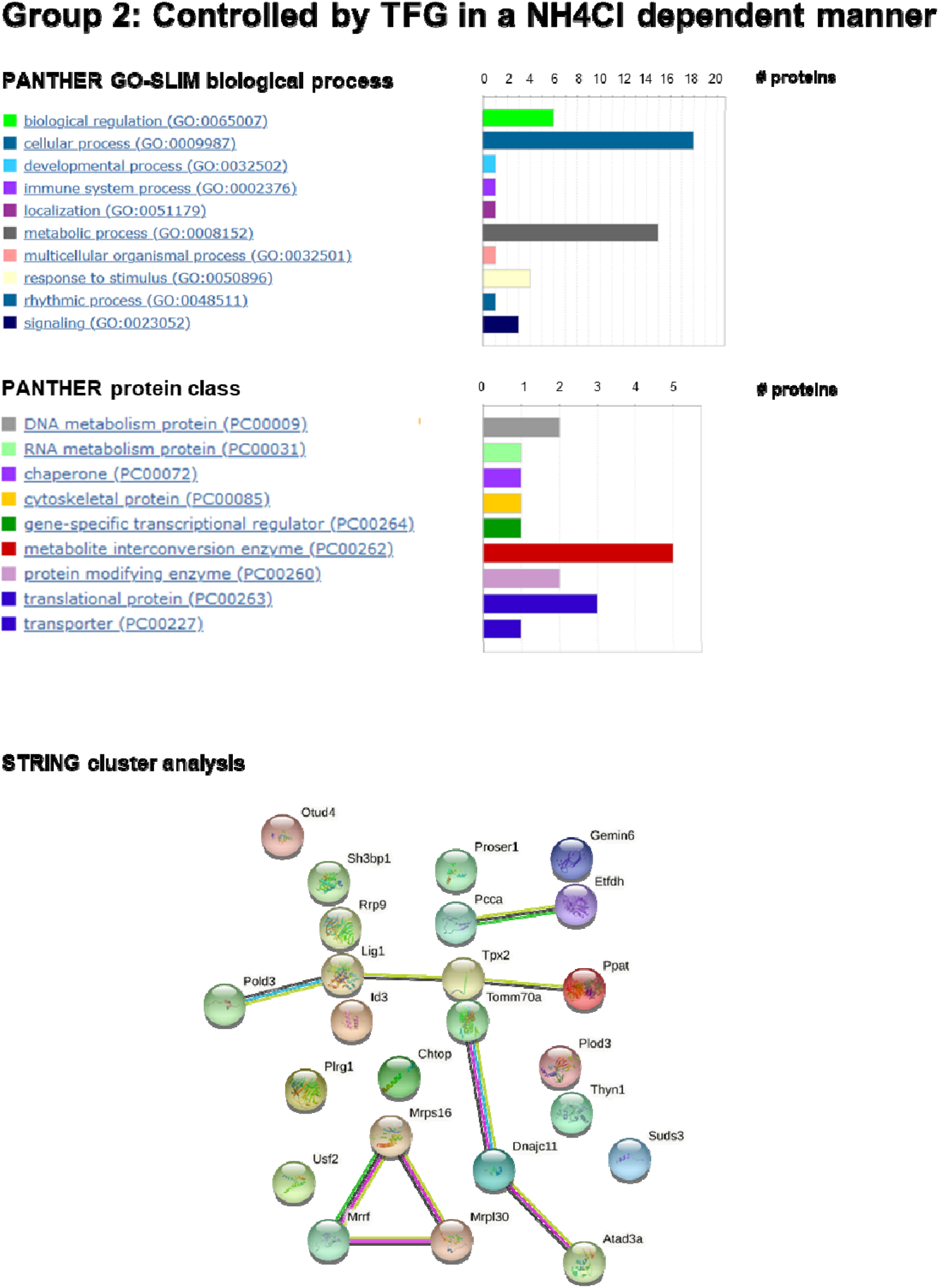
PANTHER GO-SLIM/ protein classifications as well as STRING analysis of TFG and lysosome regulated proteins. Analyses were performed with the proteins TPX2, GEMIN6, PLOD3, THYN1, MRRF, USF2, MRPS16, RRP9, CHTOP, PLRG1, OTUD4, TOMM70A, PCCA, DNAJC11, MRPL30, PPAT, POLD3, ETFDH, LIG1, ATAD3, SUDS3, PROSER1, SH3BP1, ID3

**Figure S3.**
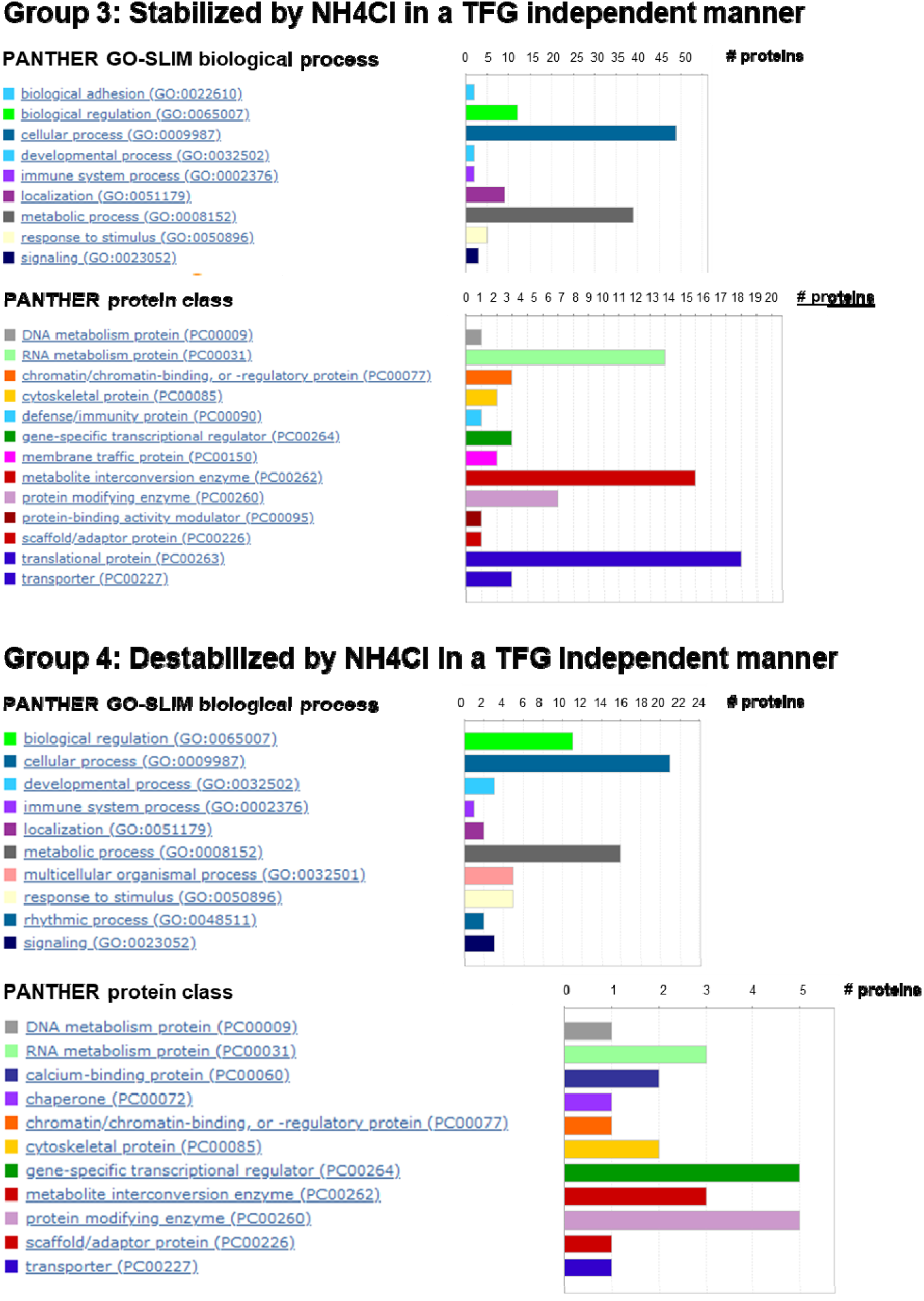
PANTHER GO-SLIM/ protein classifications of lysosome regulated proteins. Analyses were performed with the proteins RPL35A, RPL34, MRPL27, ELAC2, MTA2, BLVRA, GTPBP1, WDR43, CENPV, RPS11, ZNF622, ZC3H15, SCYL1, DBR1, EIF2B4,PFKM, UQCR10, TOMM20, DUS3L, EEFSEC, B2M, RCL1, SUZ12, MRPL3, FAM98B, WDR11, MRPL16, GPS1, H2AFV, BRD2, H2AFX, RPS2, MRPS2, RPS16, ILK, TRMT1, SLC2A3, SLC25A11, GMDS, TARBP2, GNPDA2, CWF19L1, ACP6, FARSA, PRPSAP2, TEX30, RPS25, ERLEC1, DDX1, THUMPD1, MRPL45, IFI35, HIST1H2BC, MAPK3, SCFD2, FARSB, SND1, ZNF326, EPB41, POLR1E, GPI, CHTF18, ALDH18A1, ARPC1B, ACLY, NDUFV1, OGT, RASAL1, UTP18, HDLBP, CDC73, ILF3, MIOS, FBL, PGAM5, PITRM1, PGAM2, NSF, RPS5, SRP14, UQCRQ, WARS (group 3) and ZADH2, SUDS3, ACBD6, LRRFIP2, TSC22D4, PROSER1, DNA2, TRP53, NCSTN, 5730455P16RIK, UBE2S, CCDC101, MRM1, ACADSB, N6AMT2, SH3BP1, DHPS, ECHDC2, TAF7, BOD1L, TXNIP, CHAF1A, CCDC85C, HMG20A, RRBP1, BHLHE41, ACBD3, PDE4DIP, RUNX3, UBE2B, GIGYF2, YLPM1, CCDC86, SLC7A1, NUCB1, MAD2L1, TPM3-RS7, CCT6B, ID3, SDF4, SCPEP1, USP48, TPM3 (group 4).

**Figure S4.**
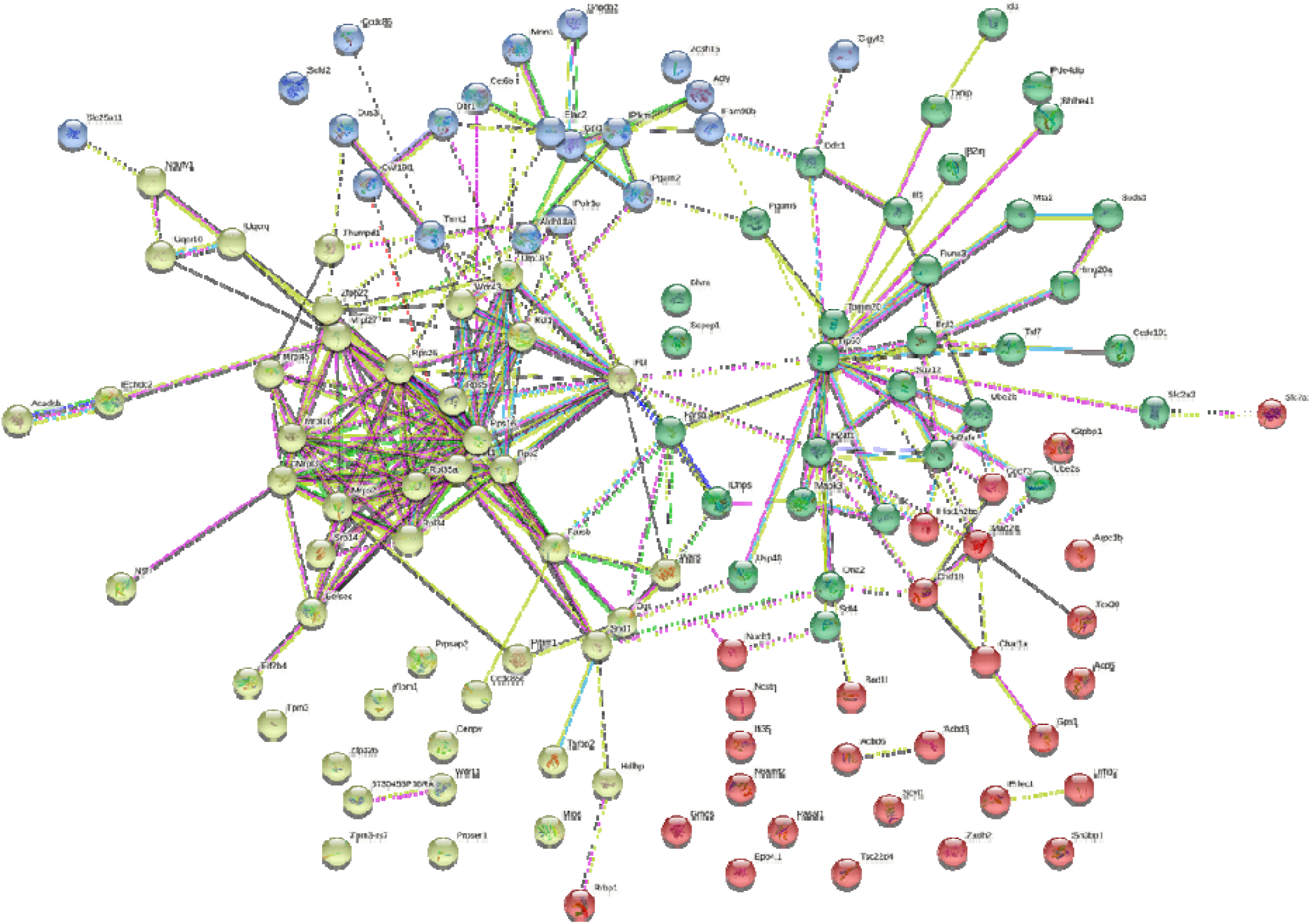
STRING analysis of lysosome regulated proteins. STRING analyses centered around 3 nodes (k-means 3) was performed with the proteins RPL35A, RPL34, MRPL27, ELAC2, MTA2, BLVRA, GTPBP1, WDR43, CENPV, RPS11, ZNF622, ZC3H15, SCYL1, DBR1, EIF2B4,PFKM, UQCR10, TOMM20, DUS3L, EEFSEC, B2M, RCL1, SUZ12, MRPL3, FAM98B, WDR11, MRPL16, GPS1, H2AFV, BRD2, H2AFX, RPS2, MRPS2, RPS16, ILK, TRMT1, SLC2A3, SLC25A11, GMDS, TARBP2, GNPDA2, CWF19L1, ACP6, FARSA, PRPSAP2, TEX30, RPS25, ERLEC1, DDX1, THUMPD1, MRPL45, IFI35, HIST1H2BC, MAPK3, SCFD2, FARSB, SND1, ZNF326, EPB41, POLR1E, GPI, CHTF18, ALDH18A1, ARPC1B, ACLY, NDUFV1, OGT, RASAL1, UTP18, HDLBP, CDC73, ILF3, MIOS, FBL, PGAM5, PITRM1, PGAM2, NSF, RPS5, SRP14, UQCRQ, WARS (group 3) and ZADH2, SUDS3, ACBD6, LRRFIP2, TSC22D4, PROSER1, DNA2, TRP53, NCSTN, 5730455P16RIK, UBE2S, CCDC101, MRM1, ACADSB, N6AMT2, SH3BP1, DHPS, ECHDC2, TAF7, BOD1L, TXNIP, CHAF1A, CCDC85C, HMG20A, RRBP1, BHLHE41, ACBD3, PDE4DIP, RUNX3, UBE2B, GIGYF2, YLPM1, CCDC86, SLC7A1, NUCB1, MAD2L1, TPM3-RS7, CCT6B, ID3, SDF4, SCPEP1, USP48, TPM3 (group 4).

**Figure S5.**
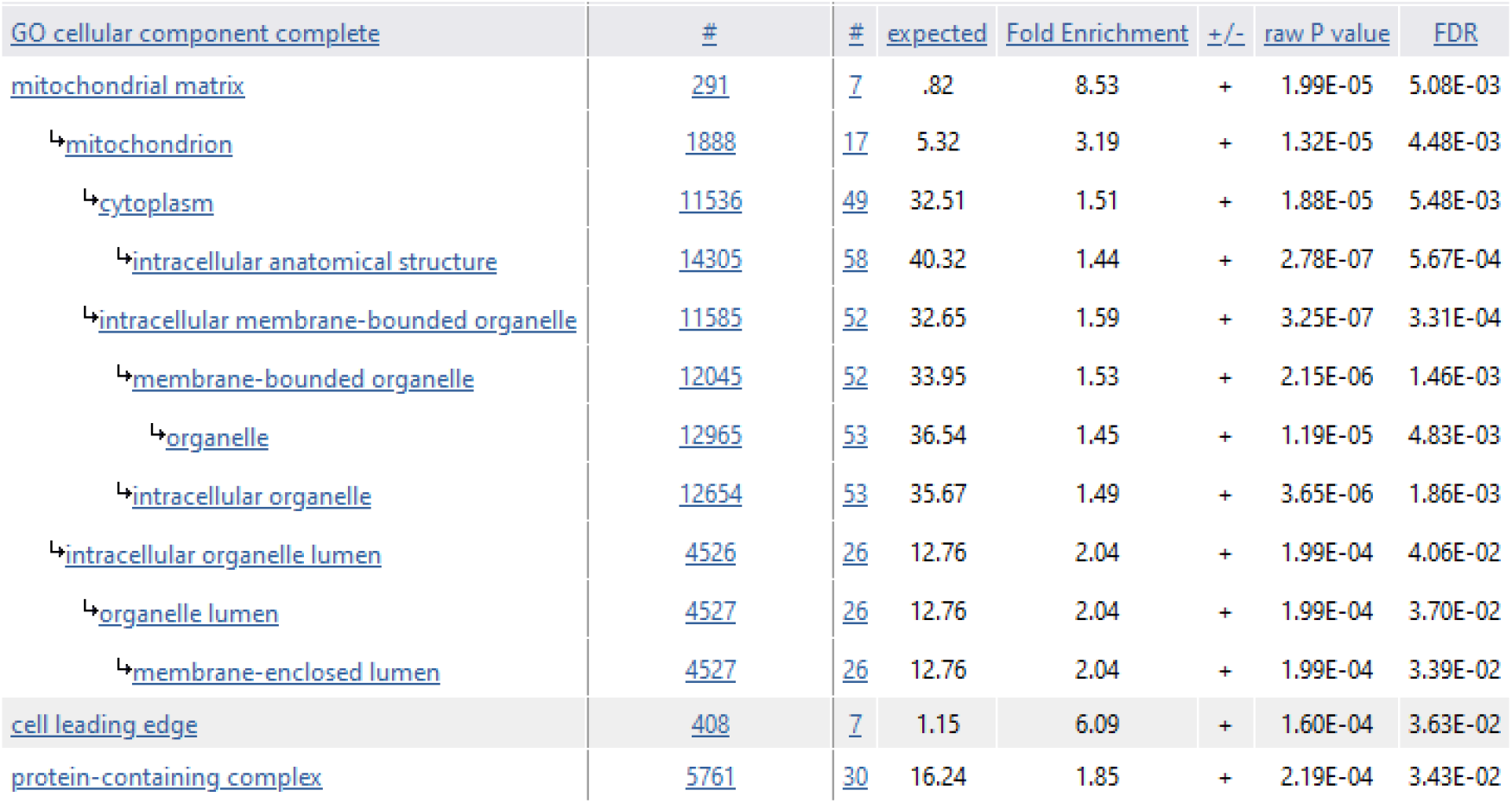
PANTHER protein overrepresentation analysis of TFG regulated proteins. Analysis was performed with the proteins IFI205A, CD36, CPNE2, IGJ, ACOT9, KNSTRN, SNW1, HSDL2, KIF5B, NUBPL, AARS2, SH3KBP1, RCOR1, MPP6, SPATS2, AAK1, TOPBP1, NAB2, FHL3, PML, E2F4, CAPNS1, ARHGAP25, GSN, COG3, EPN1, BCL10, JUNB, CTH, AVIL, PLEK, MPST, RNF113A2, CD44, PRMT6, IMPDH1, ALDOC, TPX2, GEMIN6, PLOD3, THYN1, MRRF, USF2, MRPS16, RRP9, CHTOP, PLRG1, OTUD4, TOMM70A, PCCA, DNAJC11, MRPL30, PPAT, POLD3, ETFDH, LIG1, ATAD3, SUDS3, PROSER1, SH3BP1, ID3

